# Stress granule fusion is a mitochondria-coordinated process for stress adaptation

**DOI:** 10.64898/2026.08.06.743209

**Authors:** Tae Lim Park, Geunhee Kim, Hyeong-In Kim, Sanghyun Do, Kwangmin Ryu, Yig Ji Lee, Chiyeol Song, Hwanyong Shim, Dong-Kyun Kim, Yoon Ki Kim, Won-Ki Cho

## Abstract

Stress granules are cytoplasmic membraneless organelles assembled during stress to maintain cellular homeostasis. Although fusion is a hallmark of liquid-like behavior of these condensates, whether this process carries functional significance beyond its physical coalescence remains unclear. Here, we show that stress granule fusion is facilitated by mitochondrial dynamics and membrane potential. Intact mitochondria actively associate with stress granules, facilitating fusion and maturation. In contrast, loss of mitochondrial membrane potential, along with disrupted mitochondrial structure or motility, weakens these interactions and reduces fusion frequency. We find that impaired fusion leads to the accumulation of immature granules that retain fewer sequestered components, which correlates with premature cell death. Remarkably, restoring mitochondrial membrane potential rescues granule fusion and enlargement, and is accompanied by increased cell viability and a corresponding increase in granule-associated apoptotic factors. These results demonstrate that stress granule fusion is actively coordinated by mitochondria rather than driven solely by passive coalescence, reshaping how condensate dynamics are understood to integrate with organelle function during cellular stress response.

## Introduction

Acute stress triggers the transient formation of stress granules via liquid–liquid phase separation (Protter and Parker, 2016; Lin et al, 2015). By sequestering translationally stalled mRNAs, RNA-binding proteins, and apoptotic factors (Youn et al, 2019), stress granules mitigate stress-induced damage and promote cell survival under harsh environments (Kedersha and Anderson, 2002; Anderson and Kedersha, 2008). These condensates nucleate as small structures that subsequently mature into larger assemblies through coalescence (Wheeler et al, 2016; Guillén-Boixet et al, 2020; Hu et al, 2023). Their dynamic properties, including assembly, disassembly, and mobility, are tightly regulated and critical for cellular adaptation and homeostasis (Mahboubi and Stochaj, 2017; Gwon et al, 2021; Ivanov et al, 2003).

Interactions with cytoskeletal filaments and membrane-bound organelles, such as lysosomes and the endoplasmic reticulum (ER), coordinate the dynamic behaviors of stress granules (Liao et al, 2019; Lee et al, 2020; Nadezhdina et al, 2010; Chernov et al, 2009). These interorganelle interactions facilitate RNA transport, granule mobility, and clearance, indicating the importance of organelle–granule associations for coordinating efficient cellular responses. Dysregulation of stress granule dynamics has been implicated in neurodegenerative disorders, including amyotrophic lateral sclerosis (ALS) and frontotemporal dementia (FTD) (Mateju et al, 2017; Li et al, 2013; Mackenzie et al, 2017), yet how interorganelle interactions contribute to stress granule dysfunction remains incompletely understood.

Mitochondria, dynamic organelles central to energy production, redox balance, and apoptotic signaling (Zhao et al, 2002), have recently emerged as key nodes in interorganelle coordination of stress granule dynamics. Through fission, fusion, transport, and changes in mitochondrial membrane potential (MMP), they continuously adapt to fluctuating cellular demands (Bereiter-Hahn and Vöth, 1994; Westermann, 2010). Given that both mitochondria and stress granules orchestrate stress responses, multiple lines of evidence indicate functional crosstalk: stress granules formed under starvation can modulate mitochondrial permeability and fatty acid oxidation (Amen and Kaganovich, 2021), whereas organelle-specific stress that activates the mitochondrial unfolded protein response (UPR^mt^) influences stress granule formation (Lopez-Nieto et al, 2025). Physical contacts between mitochondria and RNA granules have also been reported under conditions of mitochondrial oxidative stress (Ball et al, 2026), although these were defined in a distinct granule and metabolic context. These functional and physical links, together with reported genetic and pathological associations between ALS/FTD-associated mitochondrial defects and granule dysregulation (Lin and Beal, 2006; Kreiter et al, 2018; Dafinca et al, 2016), raise the question of how mitochondrial dynamics are coupled to the stress granule life cycle, including fusion and maturation.

Here we show that mitochondrial dynamics actively shape stress granule fusion and maturation. Using nanoscale imaging, we identify apposed interfaces between the two organelles and demonstrate that mitochondrial morphology, motility, and membrane potential together tune granule coalescence. Defective mitochondria fail to support fusion, yielding immature granules that remain too small to accumulate apoptotic factors efficiently, while restoring mitochondrial membrane potential rescues granule enlargement and cell survival. These findings establish mitochondria-coordinated fusion as a fundamental step in the cellular stress response.

## Results

### Nanoscale apposition between stress granules and mitochondria

The microtubule network serves as a scaffold for organelle interactions (Nadezhdina et al, 2010; Boldogh and Pon, 2007; López-Doménech et al, 2018). However, the spatial organization of stress granules and mitochondria within this network remains to be fully characterized. To investigate their spatial relationship, we conducted live-cell spinning disk confocal imaging of HeLa cells overexpressing mTagBFP2-G3BP1 and EGFP-αTub with mitochondria labeled by MitoTracker Deep Red (Fig 1A). Stress granules induced by oxidative stress upon sodium arsenite treatment appear by 15–20 min and approach a plateau by 40–60 min (Wheeler et al, 2016; Kedersha and Anderson, 2007; Li et al, 2002).

**Figure 1.**
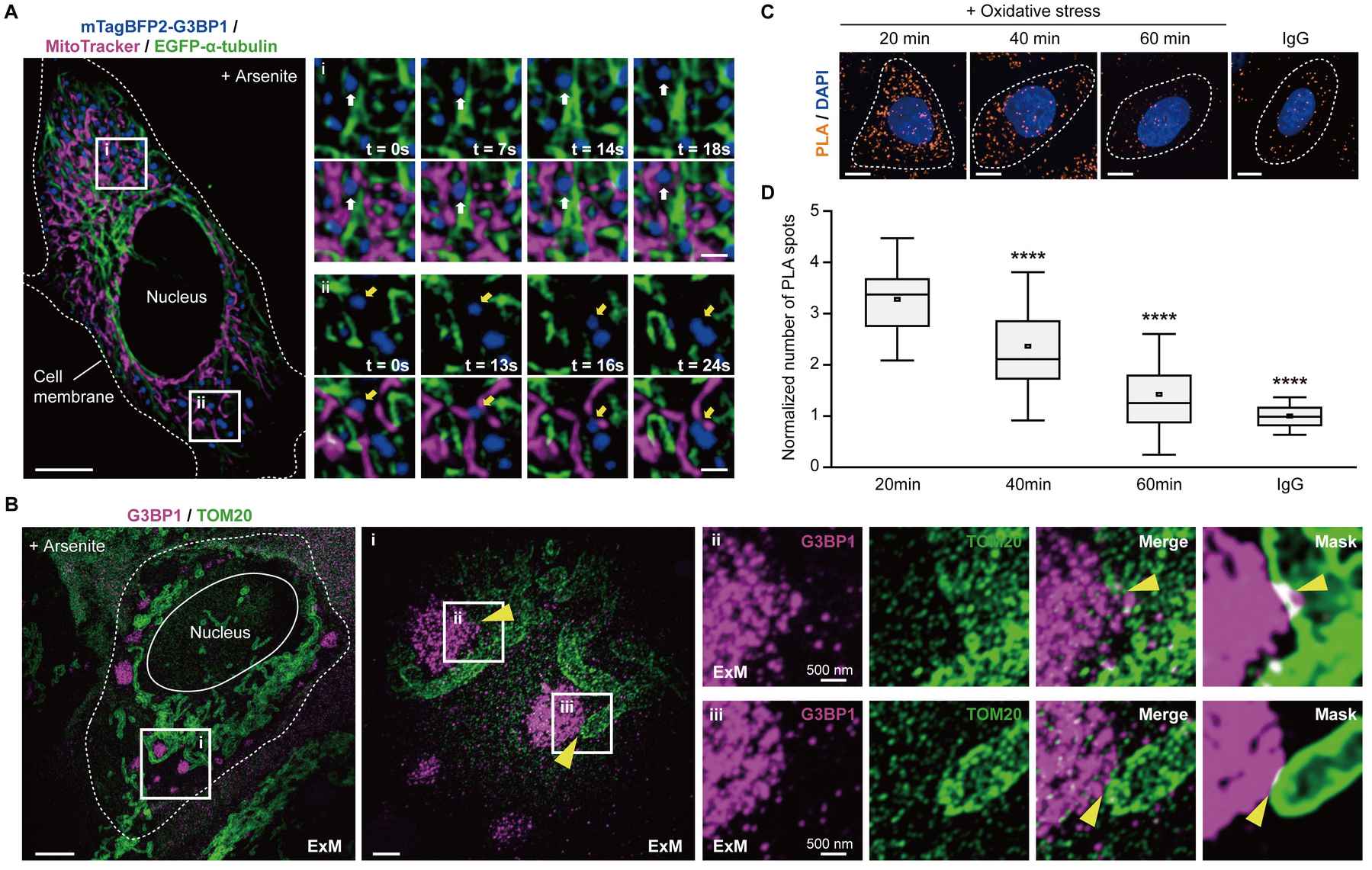
Stress granules and mitochondria form nanometer-scale spatial associations. (**A**) Representative images of live HeLa cells coexpressing mTagBFP2-G3BP1 (stress granules, blue) and EGFP-α-tubulin (microtubules, green) stained with MitoTracker Deep Red (mitochondria, magenta) after sodium arsenite treatment (500 μM, 20 min). Insets are time-lapse sequences showing the spatiotemporal distributions of stress granules and mitochondria in the presence (inset (i)) or absence (inset (ii)) of microtubules. Scale bars, 10 μm (whole cell) and 2 μm (insets). (**B**) Expansion microscopy (ExM) images of HeLa cells after sodium arsenite treatment (500 μM, 20 min) immunostained for G3BP1 (magenta) and TOM20 (green). Insets show apposed interfaces (yellow arrowheads) between stress granules and the mitochondrial outer membrane. Scale bars, 10 μm (whole cell), 2 μm (panel (i)), and 500 nm (panels (ii) and (iii)). (**C**) Representative images of PLA signals (orange) generated by anti-TOM20 and anti-G3BP1 or anti-IgG antibody pairs. Nuclei are stained with DAPI (blue). Scale bars, 10 μm. (**D**) Quantification of the PLA signals detected at 20, 40, and 60 min after 500 μM sodium arsenite treatment (*n* = 52 cells per condition). Signal intensity was normalized to the mean value of IgG group. In the box plots, center line, median; center square, mean; box limits, interquartile range; whiskers, s.d. Statistical significance was determined by one-way ANOVA with Tukey’s post-hoc test. For comparisons against the 20min group, \*\*\*\**P* < 0.0001. For IgG vs 60min, *P* = 0.0242, \**P* < 0.05.

After 20 min of arsenite exposure, stress granules exhibited two distinct spatial patterns: one population was found close to the mitochondria and microtubules (Fig 1A, inset (i)), while another subset remained adjacent to the mitochondria but distant from the microtubules (Fig 1A, inset (ii), Movie EV1). Despite this heterogeneity, both populations maintained close proximity to mitochondria during their movement. Monte Carlo simulation (Metropolis and Ulam, 1949) confirmed that stress granule–mitochondria proximity significantly exceeded values expected by chance, indicating their nonrandom spatial association (Fig EV1A, B). Given that mitochondria traffic along cytoskeletal filaments (Boldogh and Pon, 2007; López-Doménech et al, 2018), these findings raise the possibility that mitochondrial positioning may contribute to stress granule dynamics.

To determine whether stress granules and mitochondria exist in nanometer-scale apposition, we employed expansion microscopy (Pownall et al, 2023) to resolve their spatial relationship (Chen et al, 2015; Chang et al, 2017). Isotropic expansion of arsenite-treated cells followed by immunolabeling of G3BP1 and TOM20 revealed clearly apposed interfaces between stress granules and the mitochondrial outer membrane (Fig 1B, Fig EV1C). We further validated these appositions using a proximity ligation assay (PLA), which detects protein–protein proximity within 30 nm (Fig 1C, D) (Fredriksson et al, 2002). Robust PLA signals between G3BP1 and TOM20 were detected following arsenite treatment (Fig 1C), confirming their close association. Notably, these PLA signals progressively declined over time (Fig 1D), coinciding with a gradual spatial uncoupling of the two organelles during granule maturation (Fig EV1D). Together, these findings identify dynamic, nanoscale apposition between stress granules and mitochondria that diminishes as granules coalesce into larger assemblies.

### Mitochondrial dynamics facilitate stress granule fusion

Stress granules assemble through a multiple-stage process, beginning with an initial nucleation phase that forms stress granule core structures and recruits granule components, followed by a fusion-driven maturation into larger assemblies (Wheeler et al, 2016; Jain et al, 2016). Although granule fusion is a key feature of liquid-liquid phase separation (Protter and Parker, 2016; Lin et al, 2015), the mechanisms regulating this process have not been defined. Given that mitochondrial dynamics are linked to microtubule-based trafficking (Boldogh and Pon, 2007; López-Doménech et al, 2018), and microtubule disruption leads to the formation of smaller stress granules (Ivanov et al, 2003; Loschi et al, 2009), we hypothesized that mitochondria facilitate stress granule fusion during oxidative stress.

To test this, we performed live-cell imaging of HeLa cells coexpressing mNeonGreen-G3BP1 and mCherry-TOMM20 (Fig 2A). Following 20 min of arsenite exposure, we collected movies for 10 min using a cycle of 1-min imaging at 2-s intervals and 1-min rest. Mitochondria undergoing oscillatory elongation–retraction exhibited transient or semistable apposition to nearby stress granules (Fig 2A, Movie EV2). These encounters frequently coincided with stress granule fusion events, in which adjacent granules merge upon mitochondrial apposition (Fig 2A, white arrows).

**Figure 2.**
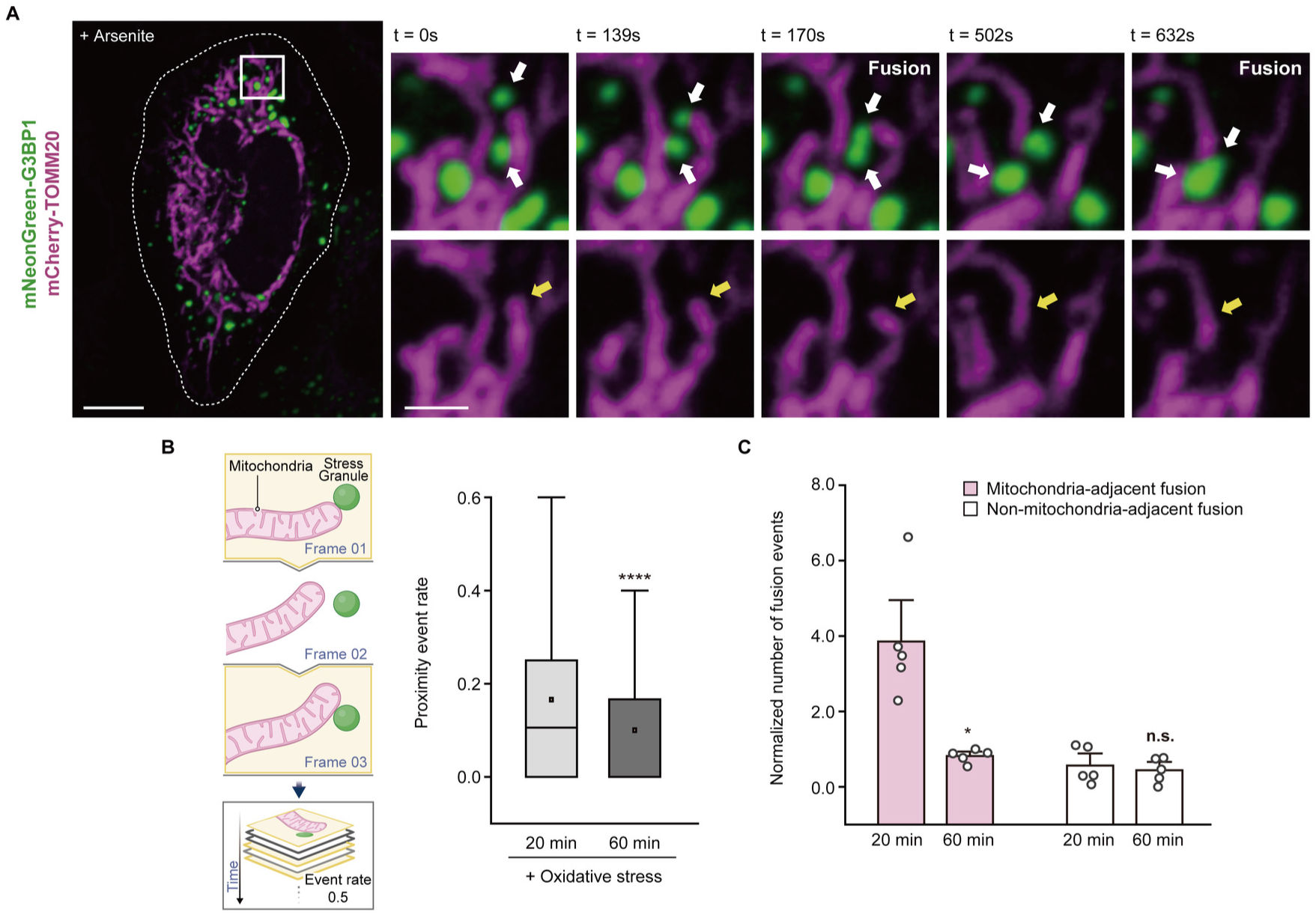
Stress granule–mitochondria appositions facilitate stress granule fusion. (**A**) Representative images of live HeLa cells expressing mNeonGreen-G3BP1 (green) and mCherry-TOMM20 (magenta) after sodium arsenite treatment (500 μM, 20 min). Insets are time-lapse images illustrating mitochondrial elongation and retraction (lower insets; yellow arrows) coinciding with stress granule fusion events (upper insets; white arrows). Scale bars, 10 μm (whole cell) and 2 μm (insets). (**B**) Schematic (left) of the analysis pipeline and quantification (right) of interorganelle proximity event rate within pre-fusion stress granule trajectories in cells exposed to the indicated durations of 500 μM sodium arsenite (*n* > 300 stress granules from five biologically independent experiments). The proximity event rate was defined as the frequency of independent close-proximity encounters normalized to the total tracking duration. In the box plots, center line, median; center square, mean; box limits, interquartile range; whiskers, outliers. Statistical significance was determined by two-tailed *t*-test. *P* = 4.69E-11 and \*\*\*\**P* < 0.0001. (**C**) Quantification of mitochondria-adjacent or non-mitochondria-adjacent stress granule fusion events in cells exposed to the indicated durations of 500 μM sodium arsenite (*n* > 250 stress granules from five biologically independent experiments). Fusion events were normalized to total granule number per cell and subsequently normalized to the minimum value across all groups. Data represent mean ± s.e.m. Statistical significance was determined by two-tailed *t*-test. *P* = 0.0140 (20 min vs 60 min, mitochondria-adjacent) and *P* = 0.65 (20 min vs 60 min, non-mitochondria-adjacent). \**P* < 0.05 and n.s., *P* ≥ 0.05.

Oxidative stress elevates cytosolic calcium and reactive oxygen species (ROS), which impair mitochondrial motility by inhibiting mitochondrial motor-adaptor proteins (Debattisti et al, 2017; Yi et al, 2004). Consistent with this, prolonged arsenite exposure considerably decreased the magnitude of mitochondrial motion, which comprises both movement capacity and directionality (Appendix Figure S1A, B). We next assessed how this stress-induced mitochondrial hypomobility influenced the physical association with stress granules. Analysis of merging trajectories specifically during the pre-fusion stage revealed that the close proximity event rate between stress granules and mitochondria was higher at 20 min compared to 60 min post-stress induction (Fig 2B). This temporal decline coincides with our previous observation that organelle association is most pronounced at early time points (Fig 1C, D).

To determine whether this proximity contributes to granule fusion, we quantified fusion events and categorized them based on their spatial context: mitochondria-adjacent fusion and non-mitochondria-adjacent fusion (e.g., diffusion-driven). Total fusion frequency reduced over time, and, notably, mitochondria-adjacent fusion, which accounted for the majority of the fusion events, markedly decreased (Fig 2C, Appendix Figure S1C). These findings suggest that mitochondrial dynamics facilitate stress granule fusion by enhancing interorganelle encounters, and that the stress-induced decline in mitochondrial motility may contribute to reduced granule coalescence and maturation during sustained stress.

### Mitochondrial dysfunction disrupts stress granule fusion and enlargement

To further investigate whether mitochondrial dynamics modulate stress granule fusion, we perturbed mitochondrial morphology and motility by depleting key regulators. Mitochondria undergo fusion, fission, and active transport to regulate their morphology, distribution, and function (Bereiter-Hahn and Vöth, 1994; Westermann, 2010). Optic atrophy 1 (OPA1) is critical for inner membrane fusion and cristae organization, and its loss leads to fragmented mitochondria (Olichon et al, 2003). Mitochondrial Rho-GTPase 1 and 2 (MIRO1/2) mediate microtubule- or actin-based mitochondrial transport, and their depletion compromises mitochondrial elongation and motility, producing shortened mitochondria with suppressed movement (López-Doménech et al, 2018; Liu and Hajnóczky, 2009).

To dissect the respective roles of mitochondrial morphology and motility in stress granule fusion, we depleted OPA1 to fragment mitochondria or suppressed MIRO1/2 to inhibit mitochondrial movement and elongation (Fig 3A, Fig EV2A–D). We acquired 10-min movies of live HeLa cells coexpressing mNeonGreen-G3BP1 and mCherry-TOMM20 after 20 min of oxidative stress, a time point when fusion events were most frequent (Fig 2C). Importantly, at this time point granule size did not differ between Si-Ctrl, Si-OPA1 and Si-MIRO1/2 cells (Fig EV2E), indicating that knockdown does not detectably alter granule nucleation and that the size differences observed at 60 min arise during the subsequent fusion-driven phase. Both perturbations significantly diminished the stress granule–mitochondria close proximity event rate (Fig 3B, Movie EV3). Consistent with this, mitochondria-adjacent fusion events markedly declined in OPA1-depleted cells and exhibited a further reduction in MIRO1/2-depleted cells (Fig 3C, Fig EV2F). Non-mitochondria-adjacent fusion was also reduced in MIRO1/2-depleted cells (Fig 3C), consistent with the broader effect of MIRO1/2 loss on cytoskeleton-dependent granule mobility.

**Figure 3.**
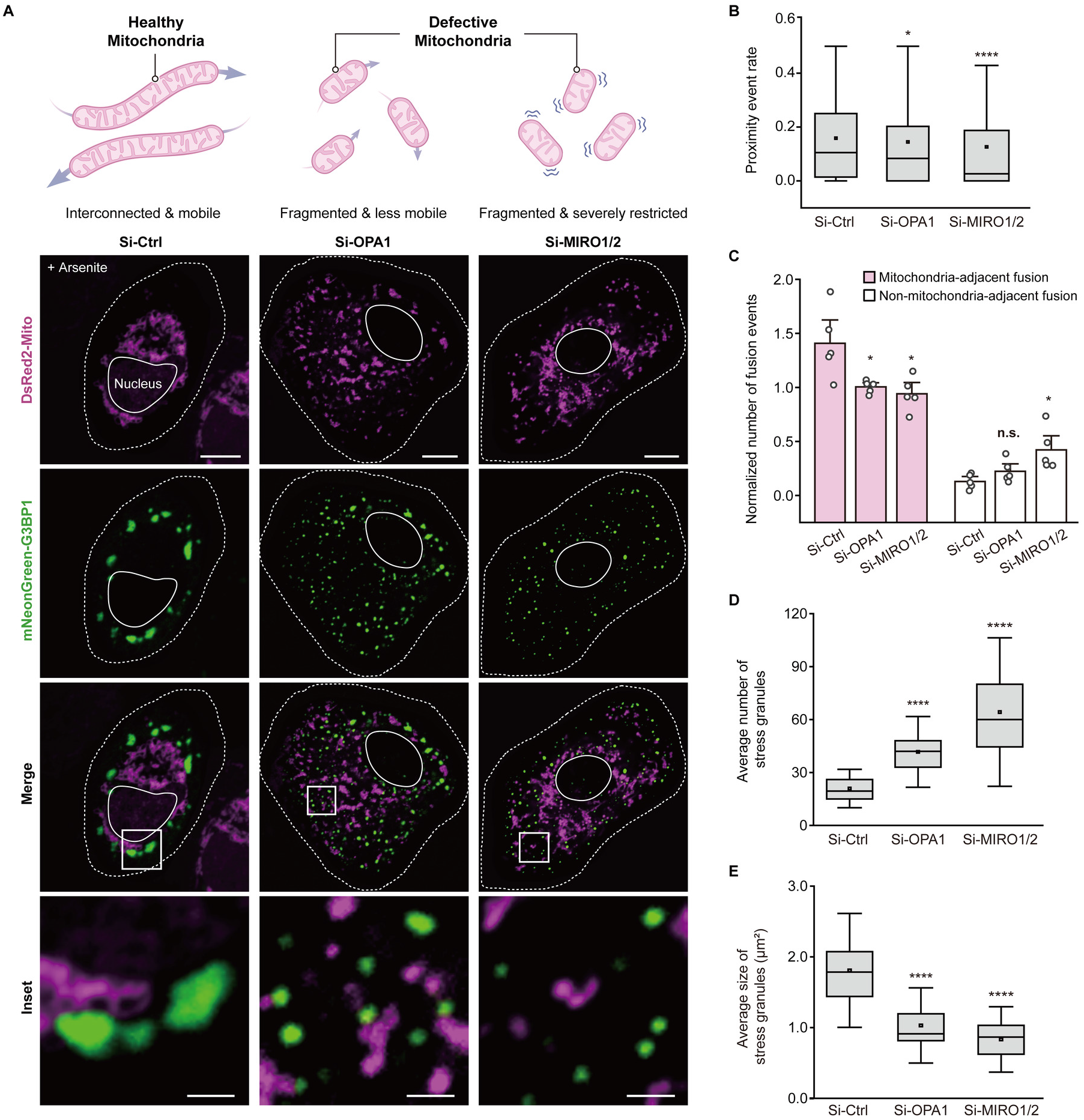
Disruption of mitochondrial morphology and motility impairs stress granule– mitochondria appositions and fusion. (**A**) Schematic (top) depicting intact versus defective mitochondria. Representative images (bottom) of Si-Ctrl, Si-OPA1, and Si-MIRO1/2 HeLa cells overexpressing DsRed2-Mito (magenta) and mNeonGreen-G3BP1 (green) after 60 min of 500 μM sodium arsenite. Scale bars, 10 μm (whole cell) and 2 μm (insets). (**B**) Quantification of interorganelle proximity event rate within pre-fusion stress granule trajectories in live HeLa cells after 20 min of 500 μM sodium arsenite treatment (*n* > 1,500 stress granules from five biologically independent experiments). In the box plots, center line, median; center square, mean; box limits, interquartile range; whiskers, outliers. Statistical significance was determined by two-tailed *t*-test. *P* = 0.0256 (Si-Ctrl vs Si-OPA1) and *P* = 5.4E-8 (Si-Ctrl vs Si-MIRO1/2). \**P* < 0.05 and \*\*\*\**P* < 0.0001. (**C**) Quantification of mitochondria-adjacent or non-mitochondria-adjacent fusion events in Si-Ctrl, Si-OPA1, and Si-MIRO1/2 HeLa cells after 20 min of 500 μM sodium arsenite treatment (*n* > 600 stress granules from five biologically independent experiments). Fusion events were normalized to total granule number per cell and then to the minimum value across all groups. Data represent mean ± s.e.m. Statistical significance was determined by two-tailed *t*-test. For mitochondria-adjacent fusion, *P* = 0.0478 (Si-Ctrl vs Si-OPA1) and *P* = 0.0280 (Si-Ctrl vs Si-MIRO1/2). For non-mitochondria-adjacent fusion, *P* = 0.1336 (Si-Ctrl vs Si-OPA1) and *P* = 0.0252 (Si-Ctrl vs Si-MIRO1/2). \**P* < 0.05 and n.s., *P* ≥ 0.05. (**D**, **E**) Quantifications of stress granule number (**D**) and size (**E**) in Si-Ctrl, Si-OPA1, and Si-MIRO1/2 HeLa cells after 60 min of 500 μM sodium arsenite treatment (*n* = 52 cells). In the box plots, center line, median; center square, mean; box limits, interquartile range; whiskers, s.d. Statistical significance was determined by two-tailed *t*-test. In (D), *P* = 2.31E-15 (Si-Ctrl vs Si-OPA1) and *P* = 1.84E-15 (Si-Ctrl vs Si-MIRO1/2). In (E), *P* = 1.44E-13 (Si-Ctrl vs Si-OPA1) and *P* =1.75E-18 (Si-Ctrl vs Si-MIRO1/2). \*\*\*\**P* <0.0001.

After 60 min of stress, cells harboring fragmented or hypomobile mitochondria displayed numerous small granules, compared with the larger, mature granules observed in control cells (Fig 3D, E), indicating that impaired fusion leads to the formation of immature granules. The extent of stress granule maturation defect correlated with the severity of mitochondrial dysfunction. In OPA1-depleted cells, fragmented mitochondria retained residual motility, allowing occasional encounters with stress granules. In contrast, MIRO1/2 loss led to a more substantial reduction in the magnitude of mitochondrial motion, thereby more severely impairing granule fusion and growth (Fig 3B, C, Fig EV2C, D). This defect exceeded that produced by blebbistatin, which blocks myosin II–mediated contractility, and was instead phenocopied by nocodazole, which disrupts microtubule-dependent mitochondrial trafficking. Nocodazole-treated cells formed even smaller granules, consistent with its broader effects on cellular architecture and the delivery of granule components (Fig EV2G, H) (Hu et al, 2023; Ivanov et al, 2003; Chernov et al, 2009). Mitochondrial motility therefore emerges as a key factor facilitating stress granule fusion and maturation.

Importantly, OPA1 and MIRO1/2 knockdown did not alter cellular ATP levels (Fig EV2I), suggesting that immature granule formation was not due to impaired nucleation or component recruitment under ATP-limiting conditions (Jain et al, 2016). Instead, it was closely linked to reduced fusion events concomitant with dysregulated mitochondrial dynamics. Comparable impairments were observed under heat shock, glucose starvation and ER stress, and at lower arsenite concentrations in HeLa cells as well as in U2OS and HEK293T cells, supporting a conserved mechanism across stress types and cellular contexts (Appendix Figure S2A–P). Osmotic stress was a partial exception, in which granule size was reduced only upon MIRO1/2 depletion.

### MMP fluctuations parallel stress granule–mitochondria association

Mitochondrial dynamics are tightly coupled with MMP, as oxidative stress–induced ROS accumulation causes mitochondrial depolarization, which in turn induces mitochondrial fragmentation (Ni et al, 2015; Park et al, 2011; Zorov et al, 2006). We first assessed changes in MMP during stress using tetramethylrhodamine ethyl ester (TMRE). Rather than declining steadily, MMP transiently increased and peaked at 15–20 min after arsenite exposure (Fig 4A), temporally coinciding with the peak in stress granule–mitochondria proximity event rate (Fig 1C, D, Fig 2B). This suggests that an early rise in MMP may support the enhanced interorganelle association.

**Figure 4.**
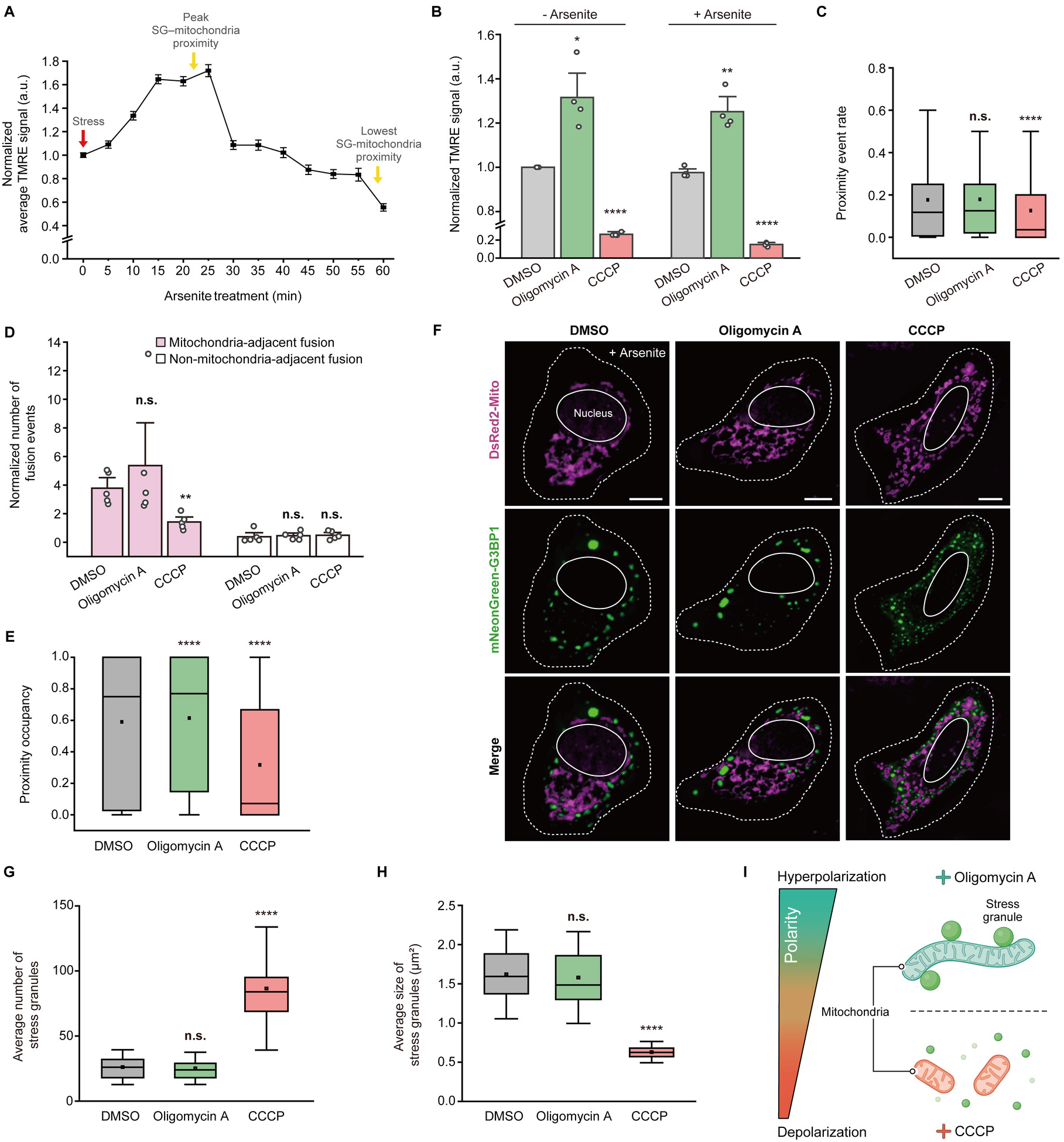
MMP modulates stress granule-mitochondria proximity occupancy and granule fusion. (**A**) Quantification of TMRE fluorescence in HeLa cells over a 60-min course of 500 μM sodium arsenite (*n* > 100 cells). Cells were imaged at 5-min intervals, and TMRE intensity at each time point was normalized to the mean intensity at 0 min. Line plots represent mean ± s.e.m. (**B**) TMRE fluorescence intensity in cells treated with the indicated chemicals before and after 500 μM sodium arsenite treatment (from 4 biologically independent experiments). Values were normalized to the DMSO-treated group. Data represent mean ± s.e.m. Statistical significance was determined by two-tailed *t*-test. At 0 min, *P* = 0.0222 (DMSO vs oligomycin A) and *P* = 4.96E-6 (DMSO vs CCCP). At 60min, *P* = 0.0054 (DMSO vs oligomycin A) and *P* = 2.94E-9 (DMSO vs CCCP). \**P* <0.05, ** *P* <0.01, and **** *P* <0.0001. (**C**) Quantification of pre-fusion stress granule–mitochondria proximity event rate in live HeLa cells treated as indicated after 20 min of 500 μM sodium arsenite treatment (*n* > 1,000 stress granules from five biologically independent experiments). In the box plots, center line, median; center square, mean; box limits, interquartile range; whiskers, outliers. Statistical significance was determined by two-tailed *t*-test. *P* = 0.3757 (DMSO vs oligomycin A) and *P* = 4.84E-15 (DMSO vs CCCP). n.s., *P* ≥ 0.05 and \*\*\*\**P* <0.0001. (**D**) Quantification of mitochondria- or non-mitochondria-adjacent fusion events in HeLa cells treated as indicated after 20 min of 500 μM sodium arsenite treatment (*n* > 300 stress granules from five biologically independent experiments). Fusion events were normalized to the total granule number per cell and then to the minimum value across all groups. Data represent mean ± s.e.m. Statistical significance was determined by two-tailed *t*-test. For mitochondria-adjacent fusion, *P* = 0.4790 (DMSO vs oligomycin A) and *P* = 0.0057 (DMSO vs CCCP). For non-mitochondria-adjacent fusion, *P* = 0.7592 (DMSO vs oligomycin A) and *P* = 0.6565 (DMSO vs CCCP). n.s., *P* ≥ 0.05, and \*\**P* < 0.01. (**E**) Quantification of the pre-fusion proximity occupancy between stress granules and mitochondria in live HeLa cells treated as indicated after 20 min of 500 μM sodium arsenite treatment (*n* > 1,000 stress granules from five biologically independent experiments). In the box plots, center line, median; center square, mean; box limits, interquartile range; whiskers, outliers. Statistical significance was determined by two-tailed *t*-test. *P* = 6.58E-6 (DMSO vs oligomycin A) and *P* = 1.14E-82 (DMSO vs CCCP). \*\*\*\**P* < 0.0001. (**F**) Representative images of HeLa cells expressing DsRed2-Mito (magenta) and mNeonGreen-G3BP1 (green) treated as indicated along with 60 min of 500 μM sodium arsenite treatment. Scale bar, 10 μm. (**G**, **H**) Quantification of the number (**G**) and size (**H**) of stress granules following 60 min of 500 μM sodium arsenite exposure in cells treated as indicated (*n* = 50 cells). In the box plots, center line, median; center square, mean; box limits, interquartile range; whiskers, s.d. Statistical significance was determined by two-tailed *t*-test. In (**G**), *P* = 0.5856 (DMSO vs oligomycin A) and *P* = 1.3E-18 (DMSO vs CCCP). In (H), *P* = 0.5975 (DMSO vs oligomycin A) and *P* = 1.16E-24 (DMSO vs CCCP). n.s., *P* ≥ 0.05, and \*\*\*\**P* < 0.0001. (**I**) Schematic model illustrating that unlike CCCP-induced depolarization, oligomycin A–induced mitochondrial hyperpolarization enhanced stress granule-mitochondria proximity occupancy.

Given the role of mitochondrial morphology and motility in facilitating stress granule fusion (Fig 3A–E), we next investigated whether MMP in defective mitochondria contributes to this process. To test this, we measured TMRE signals following OPA1 or MIRO1/2 depletion. Consistent with previous studies on mitochondrial impairments leading to depolarization (Olichon et al, 2003; Lee et al, 2022), MMP was lower in both OPA1- and MIRO1/2-deficient cells under basal conditions and further declined following arsenite exposure compared with that in intact mitochondria; this reduction was most pronounced and reached statistical significance in the MIRO1/2-deficient background (Appendix Figure S3A–C). These observations link reduced MMP in dysfunctional mitochondria to diminished encounters with stress granules.

### MMP underlies mitochondria-associated stress granule fusion by modulating the interorganelle proximity occupancy

To test the effect of MMP changes on stress granule–mitochondria association, we manipulated MMP with oligomycin A to initiate hyperpolarization and carbonyl cyanide m-chlorophenyl hydrazone (CCCP) to promote depolarization (Fig 4B, Appendix Figure S3D, E) (Billingham et al, 2022; Rainbolt et al, 2016). Mitochondrial hyperpolarization maintained stable proximity event rate between stress granules and mitochondria, whereas the depolarization yielded a significant reduction in their appositions (Fig 4C). Accordingly, mitochondria-adjacent stress granule fusion was unaffected by oligomycin A but markedly diminished by CCCP (Fig 4D, Fig EV3A), suggesting that MMP depolarization correlates with decreased interorganelle coupling.

Interestingly, despite reducing mitochondrial motility (Fig EV3B, C), oligomycin A–treated mitochondria maintained frequent apposition to stress granules (Fig 4C). Given that increased MMP has been reported to promote the colocalization of ribonucleoprotein granules with mitochondria (Cheng et al, 2022), we asked whether hyperpolarized mitochondria associated with stress granules for a larger fraction of the pre-fusion period. To investigate this, we measured the proximity occupancy of stress granules relative to mitochondria under various MMP conditions. Mitochondrial hyperpolarization significantly elevated this occupancy, whereas depolarization reduced it (Fig 4E, Movie EV4). These findings show that proximity occupancy tracks membrane potential independently of mitochondrial motility (Fig 4C–E, EV3B, C), raising the possibility that hyperpolarization could offset a motility deficit.

We next examined the effects of MMP on stress granule maturation by analyzing granule features (Fig 4F). In CCCP-treated cells, stress granules remained small and abundant, whereas in oligomycin A–treated cells, they appeared coalesced despite comparable ATP depletion in both conditions (Fig 4G, H, Fig EV3D). The accumulation of small granules upon mitochondrial depolarization indicates impaired fusion and failure to support granule enlargement. In contrast, hyperpolarized mitochondria preserved the capacity to support coalescence, demonstrating that MMP modulates stress granule maturation independently of ATP levels. Collectively, these findings show that defects in granule maturation are closely linked to dysregulated MMP rather than ATP loss. Thus, MMP, in conjunction with mitochondrial morphology and motility, modulates the interorganelle association dynamics to promote stress granule fusion and subsequent enlargement (Fig 4I).

### Mitochondrial hyperpolarization restores stress granule enlargement compromised by impaired mitochondrial dynamics

To determine whether mitochondrial hyperpolarization can rescue defective stress granule maturation caused by impaired mitochondrial dynamics, we treated OPA1- or MIRO1/2-depleted HeLa cells with oligomycin A to induce hyperpolarization (Fig EV4A–C) and monitored stress granule–mitochondria proximity dynamics following arsenite exposure. Oligomycin A treatment significantly increased interorganelle proximity occupancy in both knockdown conditions, although occupancy did not return to control level (Fig 5A). In the same recordings, the close proximity event rate, which OPA1 or MIRO1/2 depletion had reduced (Fig 3B), was no longer significantly different from that of control cells (Fig 5B). However, only OPA1-deficient cells recovered mitochondria-adjacent stress granule fusion events to control levels (Fig EV4D, E). Concurrent with this restoration, only OPA1-depleted cells restored stress granule number and size upon hyperpolarization, whereas MIRO1/2-depleted cells still accumulated immature granules (Fig 5C–E).

**Figure 5.**
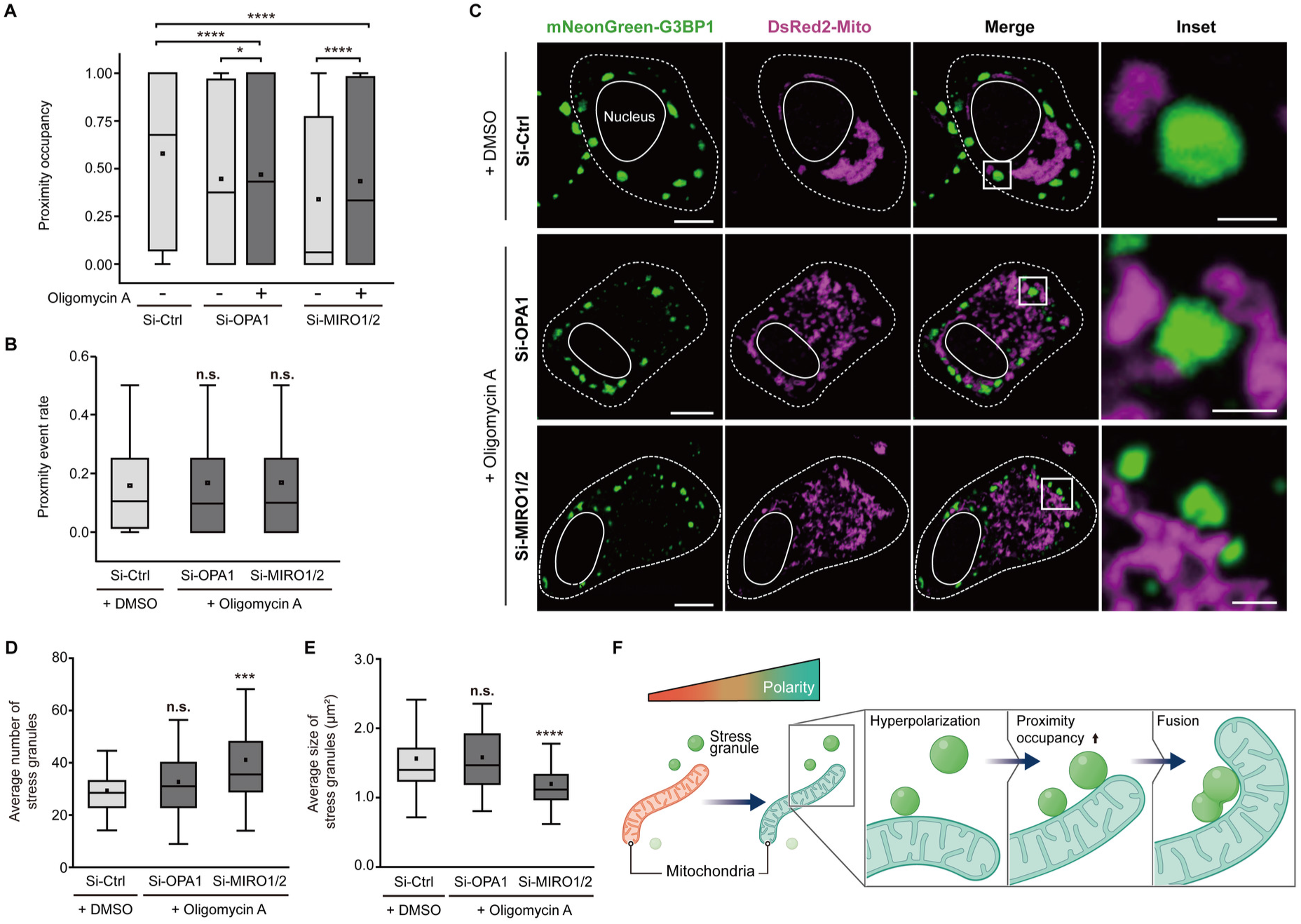
Hyperpolarization induced in defective mitochondria restores their capacity for achieving stress granule enlargement. (**A**, **B**) Quantification of pre-fusion proximity occupancy between stress granules and mitochondria (**A**) and interorganelle proximity event rate (**B**) in live HeLa cells with or without oligomycin A treatment after 20 min of 500 μM sodium arsenite treatment (*n* > 1,500 stress granules from five biologically independent experiments). In the box plots, center line, median; center square, mean; box limits, interquartile range; whiskers, outliers. Statistical significance was determined by two-tailed *t*-test. In (**A**), *P* = 6.99E-22 (Si-Ctrl vs Si-OPA1) and *P* = 2.07E-75 (Si-Ctrl vs Si-MIRO1/2). Upon oligomycin A treatment, *P* = 0.0434 (Si-OPA1 vs Si-OPA1 + oligomycin A), *P* = 4.62E-15 (Si-MIRO1/2 vs Si-MIRO1/2 + oligomycin A), *P* =7.79E-19 (Si-Ctrl vs Si-OPA1 + oligomycin A), and *P* = 5.97E-31 (Si-Ctrl vs Si-MIRO1/2 + oligomycin A). In (**B**), *P* = 0.1255 (Si-Ctrl vs Si-OPA1 + oligomycin A) and *P* = 0.0903 (Si-Ctrl vs Si-MIRO1/2 + oligomycin A). n.s., *P* ≥ 0.05, \**P* < 0.05, and \*\*\*\**P* <0.0001. (**C**) Representative images of DMSO-treated Si-Ctrl, oligomycin A–treated Si-OPA1, or oligomycin A–treated Si-MIRO1/2 HeLa cells after 60 min of 500 μM sodium arsenite treatment. Scale bars, 10 μm (whole cell) and 2 μm (insets). (**D**, **E**) Quantification of the number (**D**) and size (**E**) of stress granules following 60 min of 500 μM sodium arsenite exposure in cells treated as indicated (Si-Ctrl, *n* = 46; Si-OPA1, *n* = 49, and Si-MIRO1/2, *n* = 50 cells). In the box plots, center line, median; center square, mean; box limits, interquartile range; whiskers, s.d. Statistical significance was determined by two-tailed *t*-test. In (**D**), *P* = 0.2233 (Si-Ctrl + DMSO vs Si-OPA1 + oligomycin A) and *P* = 0.0002 (Si-Ctrl + DMSO vs Si-MIRO1/2 + oligomycin A). In (**E**), *P* = 0.8812 (Si-Ctrl + DMSO vs Si-OPA1 + oligomycin A) and *P* = 5.75E-23 (Si-Ctrl + DMSO vs Si-MIRO1/2 + oligomycin A). n.s., *P* ≥ 0.05, and \*\*\**P* <0.001, \*\*\*\**P* <0.0001. (**F**) Schematic model summarizing that the hyperpolarization of impaired mitochondria restores stress granule fusion and maturation by enhancing the association.

Because the event rates in both backgrounds were no longer distinguishable from control (Fig 5B), this divergence cannot be attributed to interorganelle apposition alone. What distinguishes the two backgrounds is residual mitochondrial motion, which oligomycin A did not restore in either case (Fig EV4F, G). MIRO1/2-depletion markedly reduced mitochondrial movement, whereas OPA1-deficient mitochondria retained relatively higher motility (Fig EV2C, D). Sustained apposition therefore appears to support fusion only when some mitochondrial motility remains, so that hyperpolarization can offset a partial but not a near-complete loss of motion. We note that MIRO1/2 also participate in ER-mitochondria contacts and mitochondrial Ca^2+^ handling, so additional motility-independent contributions of these GTPases to the rescue defect cannot be excluded. Together, these findings are consistent with a model in which mitochondrial hyperpolarization compensates for partial loss of motility by increasing interorganelle proximity occupancy, thereby facilitating stress granule fusion and enlargement (Fig 5F).

### Stress granule fusion is essential for efficient protein sequestration

Apart from their role as transient storage sites for various RNAs and proteins, stress granules also recruit proapoptotic factors such as RACK1 and Raptor (Arimoto et al, 2008; Thedieck et al, 2013) to suppress apoptotic signaling and prevent premature cell death under adverse conditions. Stress granules grow via Ostwald ripening and coalescence (Wheeler et al, 2016; Ostwald, 1900), processes that reduce surface-to-volume ratio and interfacial spaces (Welsh et al, 2022; Lee et al, 2023; Bergeron-Sandoval and Michnick, 2018; Brangwynne, 2011). Because larger granules potentially provide higher local concentrations and increased molecular interactions among resident components (Shin and Brangwynne, 2017; Brangwynne et al, 2009; Protter et al, 2018), we hypothesized that stress granule enlargement increases the efficiency of proapoptotic factor sequestration, that is, that granule content rises faster than granule volume alone would predict.

To examine the size-dependent capacity of stress granules to recruit apoptotic proteins, we performed dual-color direct stochastic optical reconstruction microscopy (dSTORM) to localize target proteins at single-molecule resolution (Heilemann et al, 2008; Heilemann et al, 2009). Cells harboring either intact or dysfunctional mitochondria were subjected to oxidative stress and analyzed for colocalization of endogenous RACK1 or overexpressed HA-mCherry–tagged Raptor with the stress granule marker G3BP1 (Fig 6A, Appendix Figure S4A). Because both the localization count and the granule area are measured within the same optical section, constant internal concentration predicts a scaling exponent of 1.5 for granules contained within that section and of 1 for granules extending beyond it; both are reported as reference values, with 1.5 the more conservative. We therefore fitted a power law to the relationship between granule area and RACK1 content in each condition. In control cells, the fitted exponent was 1.67 for RACK1, significantly greater than both 1 and 1.5 (Fig 6B, Table EV1), and steeper still for Raptor at 2.49 (Appendix Figure S4B, Table EV2). The power law was better supported than either a linear or an exponential model in all conditions except Si-MIRO1/2 + oligomycin A for Raptor, where the exponential model had a marginally lower AICc.

**Figure 6.**
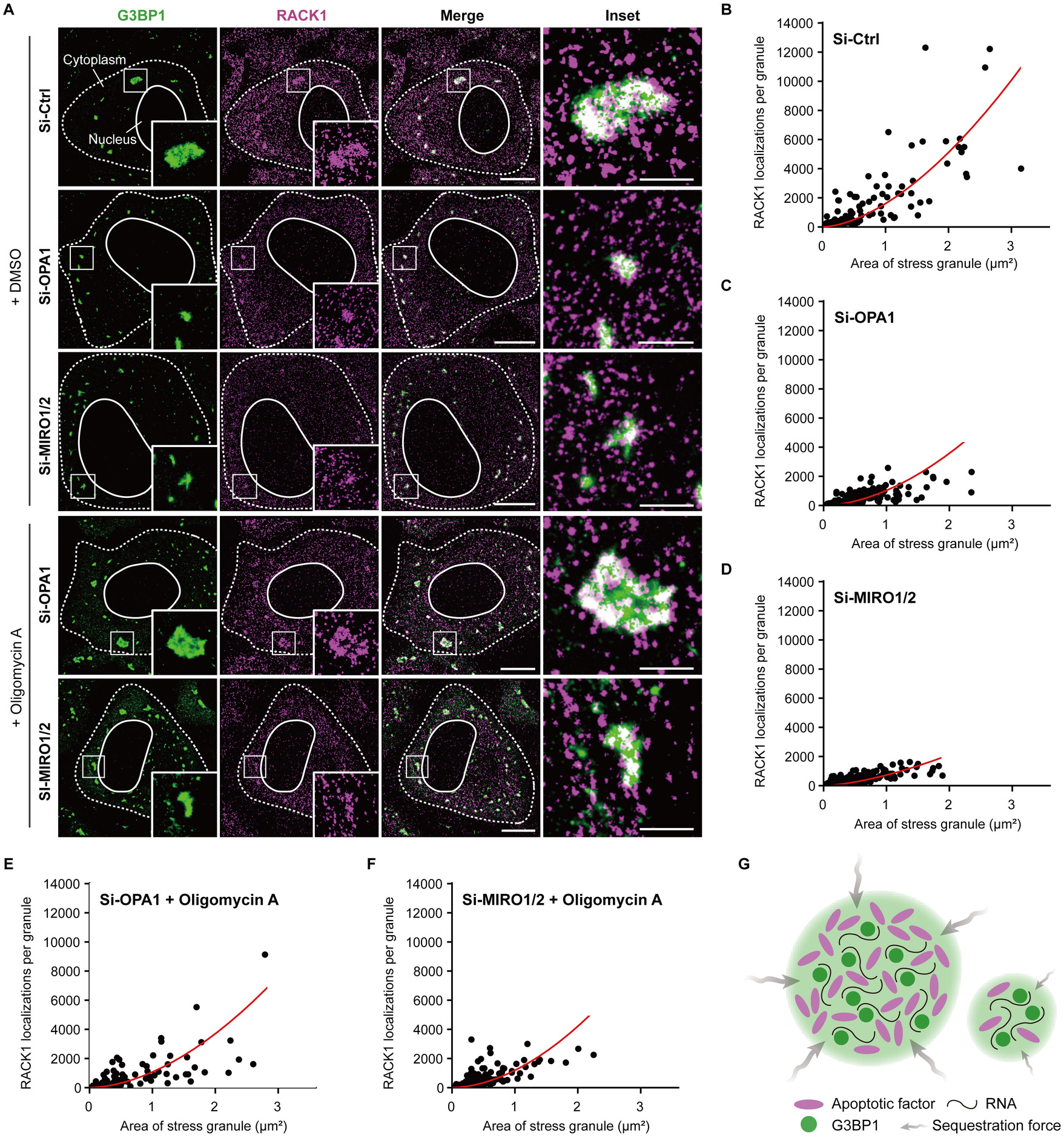
Enlarged stress granules enhance sequestration of the apoptotic factor RACK1. (**A**) Representative dual-color dSTORM images of G3BP1 (Alexa Fluor 488, green) and RACK1 (Alexa Fluor 647, magenta) in HeLa cells transfected and treated as indicated after sodium arsenite exposure (500 μM, 60 min). Scale bars, 10 μm (whole cell) and 2 μm (insets). (**B**–**F**) Number of RACK1 localizations within individual stress granules plotted against the granule area in DMSO-treated Si-Ctrl (**B**), Si-OPA1 (**C**), and Si-MIRO1/2 (**D**) cells, and oligomycin A–treated Si-OPA1 (**E**), and Si-MIRO1/2 (**F**) cells. (Si-Ctrl, n = 300; Si-OPA1, n = 539; Si-MIRO1/2, n = 718; Si-OPA1 + oligomycin A, n = 252; Si-MIRO1/2 + oligomycin A, n = 461 granules from three biologically independent experiments). Red lines denote the fitted power law y = *A*_0_ · *x*^*b*, estimated by ordinary least squares on log-transformed values. Fitted exponents, confidence intervals and model comparisons are given in Table EV1. (**G**) Schematic model illustrating that stress granule enlargement increases the amount of protein sequestered.

The exponent itself was largely preserved when mitochondrial dynamics were disrupted, differing from control in neither Si-OPA1 nor Si-MIRO1/2 cells (Fig 6C, D, Table EV1), although in Si-MIRO1/2 cells it was no longer distinguishable from 1.5. What changed instead was the range of granule sizes over which that relationship could operate (Table EV1). Because content rises steeply with area, the high-content regime is reached only by granules that grow large, and OPA1- or MIRO1/2-depleted cells rarely reach it (Fig 3E). Consistently, the RACK1 exponent under oligomycin A remained indistinguishable from control in both backgrounds (Fig 6E, F, Table EV1), yet only OPA1-depleted cells regained large granules (Fig 5E) and the corresponding gain in content. What mitochondrial dysfunction constrains is therefore not the scaling itself but the granule size that can be attained. In every condition, Raptor content also increased with granule area (Appendix Figure S4B-F), independently confirming that recruitment scales with granule size. The fitted exponent and its absolute level differed between conditions (Table EV2); because Raptor was assessed as an overexpressed HA-mCherry fusion, we base the scaling analysis on endogenous RACK1.

To further evaluate whether this size-dependent recruitment applies to other stress granule– or mitochondria–associated proteins, we quantified fluorescence intensities of eIF3η, caspase-3, VCP and proteasome 20S within G3BP1-labeled granules. All examined proteins accumulated less within granules in OPA1- or MIRO1/2-depleted cells (Appendix Figure S5A–D), consistent with the smaller granules formed under these conditions (Fig 3E). Together, these results demonstrate that granule enlargement is accompanied by greater recruitment of these proteins as well, consistent with the superlinear scaling resolved at single-molecule resolution for RACK1 and Raptor, and indicating that fusion-driven growth is functionally consequential rather than merely additive (Fig 6G).

### Large, mature stress granules attenuate apoptosis and promote cell recovery against cellular stress

Upon stress relief, stress granules disassemble through the decondensation of their components (Wheeler et al, 2016; Hofmann et al, 2021), a process critical for cell survival by enabling the resumption of cellular activities and the restoration of homeostasis. Given that fusion-driven enlargement enhances the sequestration capacity of stress granules (Fig 6), we tested whether immature granules resulting from mitochondrial dysfunction impair cell survival. To assess disassembly kinetics, we quantified the percentage of stress granule– positive HeLa cells expressing mNeonGreen-G3BP1 during recovery. Under normal conditions, stress granules resolve within 3 hours after stress relief (Wheeler et al, 2016; Gwon et al, 2021). However, granule clearance was delayed in both OPA1- and MIRO1/2-depleted cells, reaching statistical significance in the MIRO1/2-depleted background (Fig 7A, Fig EV5A). Notably, oligomycin A treatment, which restored stress granule maturation in these cells, accelerated disassembly to rates comparable to control cells (Fig 7A). Impaired disassembly was accompanied by reduced sequestration of VCP and proteasome 20S (Appendix Figure S5C, D), core components of arsenite-induced stress granule clearance machinery (Turakhiya et al, 2018), suggesting that sequestration defects in immature granules compromise cell viability.

**Figure 7.**
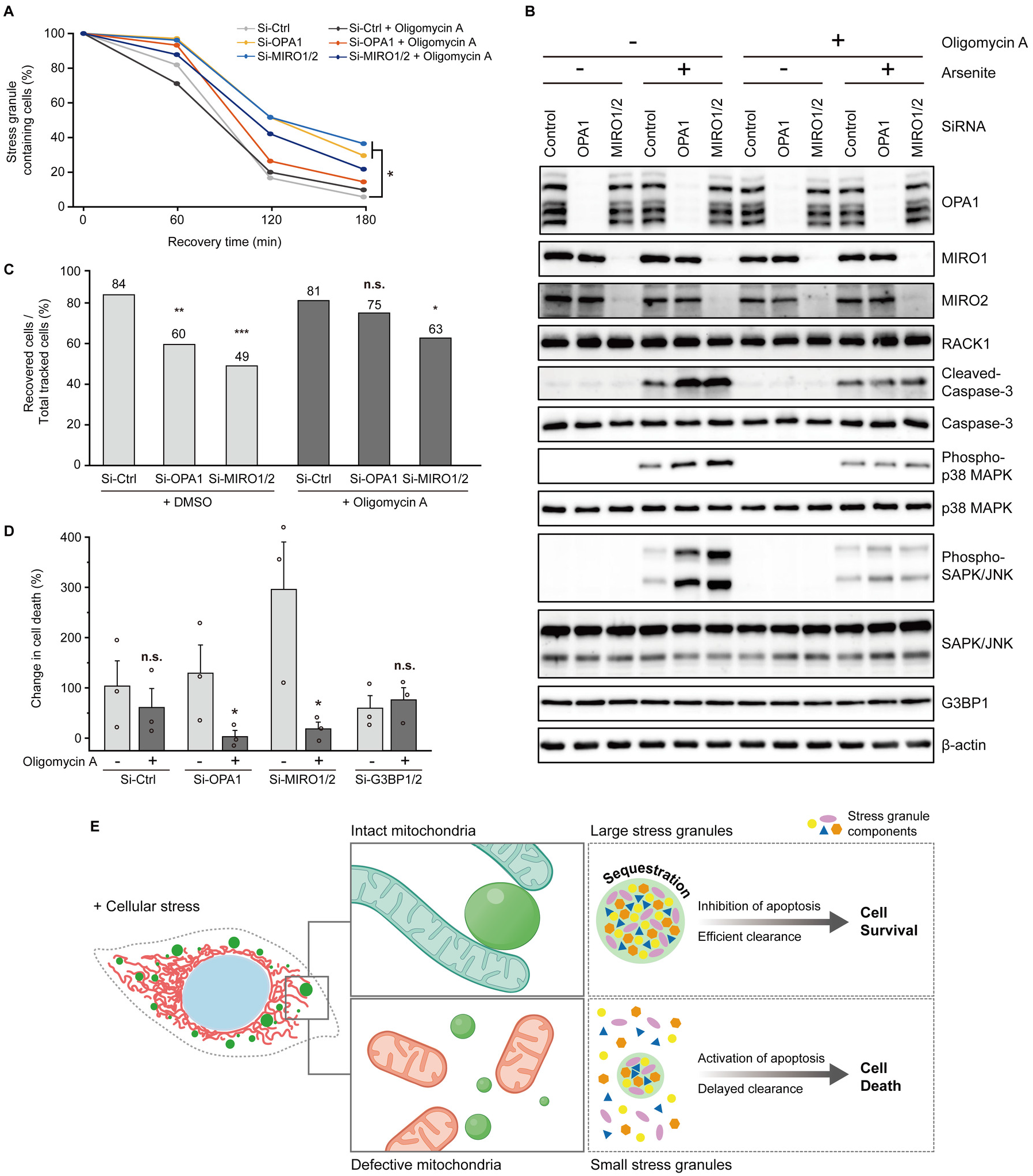
Mitochondria-dependent stress granule enlargement promotes granule clearance and cell survival. (**A**) Time-course quantification of stress granules disassembly in Si-Ctrl, Si-OPA1, and Si-MIRO1/2 HeLa cells with or without oligomycin A at 0, 60, 120, and 180 min of recovery (*n* > 79 cells per group) after 60 min of 500 μM sodium arsenite treatment. Statistical significance was determined using the Mantel–Cox test compared to Si-Ctrl group. *P* = 0.02 (Si-Ctrl vs Si-MIRO1/2) and \**P* < 0.05. (**B**) Immunoblot of apoptosis signaling molecules in control and knockdown cells treated with or without oligomycin A before and after arsenite treatment exposure (500 μM, 60 min). (**C**) Quantification of cell recovery rates based on live-cell tracking of Si-Ctrl, Si-OPA1, and Si-MIRO1/2 HeLa cells after 60 min of 500 μM sodium arsenite exposure treated with or without oligomycin A (*n* = 47-55 cells per condition). Statistical significance was analyzed using Fisher’s exact test compared to the Si-Ctrl group. *P* = 0.007 (Si-Ctrl vs Si-OPA1), *P* = 0.0002 (Si-Ctrl vs Si-MIRO1/2), *P* = 0.312 (Si-Ctrl vs Si-OPA1 + oligomycin A), and *P* = 0.021 (Si-Ctrl vs Si-MIRO1/2 + oligomycin A). \**P* < 0.05, \*\**P* < 0.01, \*\*\**P* < 0.001 and n.s., *P* ≥ 0.05. (**D**) Quantification of stress-induced apoptosis based on Annexin V staining in Si-Ctrl, Si-OPA1, Si-MIRO1/2, and Si-G3BP1/2 cells with or without oligomycin A after 60 min of 500 μM sodium arsenite treatment. Changes in cell death before and after oxidative stress were calculated from three biologically independent experiments. Data represent mean ± s.e.m. Statistical significance was determined using two-tailed *t*-test. *P* = 0.2680 (Si-Ctrl ± oligomycin A), *P* = 0.0464 (Si-OPA1 ± oligomycin A), *P* = 0.0219 (Si-MIRO1/2 ± oligomycin A), and *P* = 0.3298 (Si-G3BP1/2 ± oligomycin A). \**P* < 0.05 and n.s., *P* ≥ 0.05. (**E**) Schematic model summarizing the proposed mechanism. Under oxidative stress, intact mitochondria facilitate stress granule fusion and enlargement. Enlarged granules effectively sequester proapoptotic factors and their components, thereby increasing cell survival. In contrast, defective mitochondria fail to support granule fusion, resulting in immature granules that cannot suppress apoptosis and ultimately cause cell death.

Given the role of stress granules in sequestering proapoptotic proteins such as RACK1 (Arimoto et al, 2008), we next examined whether defective granule enlargement exacerbates apoptotic signaling. Upon arsenite exposure, OPA1- and MIRO1/2-deficient cells exhibited elevated cleaved caspase-3 expression and p38 MAPK and SAPK/JNK phosphorylation (Fig 7B), indicative of heightened apoptotic activity. Oligomycin A treatment reduced cleaved caspase-3 accumulation and phosphorylated protein levels (Fig 7B), suggesting that restored granule maturation rescues the anti-apoptotic function of stress granules. These findings support the functional importance of granule enlargement in suppressing apoptotic signaling.

To further confirm whether stress granule maturation promotes cell survival, we monitored the recovery of live HeLa cells following stress removal. Recovery through granule disassembly fell from 84% in control cells to 60% and 49% upon OPA1 and MIRO1/2 depletion, and oligomycin A restored it substantially in the OPA1 background and partially in the MIRO1/2 background, while leaving control cells unaffected (Fig 7C). Annexin V staining gave the same ordering: stress-induced cell death rose about threefold above control upon MIRO1/2 depletion, and oligomycin A suppressed this increase in both knockdown backgrounds but not in control cells (Fig 7D).

To rule out the possibility that oligomycin A promotes survival through secondary, stress granule-independent mitochondrial pathways, we evaluated its effect in G3BP1/2-depleted cells. Critically, the protective effect of oligomycin A was abolished when stress granule formation was blocked by G3BP1/2 depletion (Fig 7D, Fig EV5B–D) (Matsuki et al, 2013), indicating that the observed enhancement in cell survival is not a granule-independent mitochondrial effect. Collectively, these findings establish that optimal mitochondrial dynamics facilitate granule fusion and enlargement, thereby enhancing protein sequestration, restraining stress-induced cell death, and promoting recovery from cellular stress (Fig 7E).

## Discussion

Our study identifies mitochondria as key regulators of stress granule dynamics, revealing a function beyond their established roles in cellular metabolism. While cytoskeletal filaments are often considered the primary drivers of stress granule mobility (Liao et al, 2019; Lee et al, 2020; Nadezhdina et al, 2010), our findings suggest that mitochondria, which traffic along these networks (Boldogh and Pon, 2007; López-Doménech et al, 2018), serve as dynamic platforms that facilitate stress granule fusion and enlargement. This role positions mitochondria as active organizers of phase-separated compartments and integral components of the cellular stress response.

Stress granule formation is widely described as a spontaneous process driven by liquid–liquid phase separation (Protter and Parker, 2016; Lin et al, 2015), with fusion generally assumed to reflect passive coalescence of condensates. However, our data indicate that stress granule fusion is not solely a passive process, but is modulated by mitochondrial activity, expanding the view that fusion is governed by biophysical minimization of interfacial energy (Welsh et al, 2022; Lee et al, 2023; Bergeron-Sandoval and Michnick, 2018; Brangwynne, 2011). This framework parallels emerging concepts in nucleolar remodeling and chromatin condensates, where active factors modulate the dynamics of condensate coarsening and composition (Lafontaine et al, 2021; Lee et al, 2021).

Mitochondrial dynamics promote stress granule fusion by increasing interorganelle apposition, thereby enlarging assemblies. This size phenotype generalizes across diverse stress paradigms and cellular backgrounds. By contrast, osmotic stress behaves as a partial exception in which granule size remains unchanged between control and OPA1-depleted conditions. Prior reports that hypertonic stress fragments mitochondrial networks and compromises MMP (Copp et al, 2005) raise the possibility that sorbitol may impose mitochondrial fragmentation and low MMP in control cells, resembling OPA1 knockdown and thereby limiting apposition-driven granule growth in our system.

Functionally, the mitochondria-dependent fusion step enhances granule capacity. Larger, mature granules sequester RACK1 disproportionately to their size, so that a modest increase in granule area yields a substantially larger increase in granule content. Because this scaling is preserved even in cells with impaired mitochondrial dynamics, the principal limiting step is not the partitioning capacity of a granule but the size it is able to reach. These granules may function as protective compartments, limiting the diffusion of death-associated proteins and thereby preserving cell viability. In contrast, defects in mitochondrial morphology, motility, or membrane potential give rise to immature granules that remain too small to accumulate these factors efficiently, thereby exacerbating apoptotic signaling. These findings establish stress granule fusion as an essential determinant of cellular resilience, directly linking mitochondrial dynamics to the functional remodeling of stress-induced condensates.

A further implication concerns how condensate size relates to condensate function. Size is usually treated as a readout of assembly, but our measurements indicate that it is itself a functional variable. Because RACK1 content rises with granule area faster than proportionally, a granule that fails to grow is disproportionately impaired in what it can hold. If superlinear loading is a general property of condensates that concentrate client proteins, then any process that limits coarsening such as the mechanics of the surrounding cytoplasm (Lee et al, 2021) would carry functional consequences out of proportion to its effect on size alone.

Several limitations of our study point to important directions for further investigation. First, although our data establish that the mitochondrial membrane potential (MMP), morphology, and motility together govern stress granule-mitochondria apposition and fusion, the molecular identity of the interface remains to be defined. Mitochondria may anchor stress granules via specific protein tethers, or act as localized signaling hubs that spatially bias fusion. A candidate tether has recently been described: TDP-43 links RNA granules to mitochondria through GADD34 at the outer mitochondrial membrane (Ball et al, 2026). That mechanism was defined for TDP-43-positive granules in galactose-based OXPHOS medium, however, and whether an equivalent tether operates at the arsenite-induced appositions resolved here cannot be assumed. Second, how MMP biases interorganelle proximity is not resolved by our experiments; possibilities include MMP-dependent changes in ion gradients or redox state. The same study reported a redox-based mechanism in which mitochondrial ROS oxidizes TDP-43 to control untethering of these contacts. There, contacts increased under conditions that lower membrane potential, opposite in sign to our observations. That study, however, reported that this pathway is not engaged by sodium arsenite, the stressor used throughout our experiments, so the two sets of observations need not report on the same regulatory input. Redox- and potential-dependent inputs are therefore likely separable, and uncoupling them will be needed to define their relative contributions.

By elucidating a mitochondria-driven mechanism for stress granule enlargement, our study proposes a regulatory axis through which mitochondrial activity shapes the architecture and function of biomolecular condensates. This axis underscores the broader importance of mitochondrial homeostasis in maintaining stress granule competence and adaptive cellular responses. Mitochondrial dysfunction and aberrant stress granule dynamics co-occur in several neurodegenerative conditions, including ALS and FTD, where transport deficits and persistent, poorly resolving granules have both been reported in patient-derived neurons (Kreiter et al, 2018; Dafinca et al, 2016). Our findings raise the possibility that these two features are sequentially linked, with defective mitochondria giving rise to immature granules. This predicts that restoring mitochondrial motility or membrane potential should improve granule resolution without acting on granule components directly. Direct tests in disease-relevant models, measuring granule size and clearance alongside mitochondrial parameters, will be an important next step.

## Materials and Methods

All antibodies, chemicals, siRNAs, commercial assay kits, and software used in this study are listed in the Reagents and Tools Table (Table EV3).

### Cell culture and chemical treatments

HeLa, U2OS, and HEK293T cells (ATCC) were cultured at 37°C with 5% CO_2_ in Dulbecco’s modified Eagle’s medium (DMEM; Gibco) supplemented with 10% fetal bovine serum (FBS; Gibco) and 1% penicillin/streptomycin (Gibco). Prior to experiments, the cells were tested for Mycoplasma contamination using MycoStrip (InvivoGen).

To induce oxidative stress, cells were treated with 500 μM sodium arsenite (Sigma-Aldrich) for 1 hour unless stated otherwise. For heat shock, cells were incubated at 43°C in a humidified chamber with 5% CO_2_. For glucose starvation, cells were cultured in glucose-free DMEM (Gibco) supplemented with 10% dialyzed FBS (Gibco) and 1% penicillin/streptomycin (Gibco). For ER stress, cells were treated with 1 μM thapsigargin (Sigma-Aldrich) for 90 min. For osmotic stress, cells were treated with 0.4 M sorbitol (Sigma-Aldrich) for 1 hour. Mitochondria were labeled with 20 nM MitoTracker Deep Red (Invitrogen) for 20 min prior to arsenite treatment. To manipulate MMP, cells were treated with 50 nM oligomycin A (Sigma-Aldrich) for 30 min to induce hyperpolarization or with 100 μM CCCP (Sigma-Aldrich) for 30 min to induce depolarization. For cytoskeletal perturbation, cells were incubated with 20 μM nocodazole (Sigma-Aldrich) for 30 min to depolymerize microtubules and with 100 μM blebbistatin (Sigma-Aldrich) for 3 hours to inhibit myosin II activity.

### DNA plasmids and transfection

The following plasmids were purchased from Addgene: EGFP-Tubulin-6 (#56450), mCherry-TOMM20-N-10 (#55146), pCS2+mNeonGreen-C Cloning Vector (#128144), DsRed2-Mito-7 (#55838), pRK5-HA-mCherry-raptor (#73386), and GW1-PercevalHR (#49082). mNeonGreen-G3BP1 was constructed by cloning G3BP1 into the HindIII/BamHI sites of the mNeonGreen-C Cloning vector, followed by subcloning with mTagBFP2 to generate mTagBFP2-G3BP1.

Transfections were performed using FuGENE HD (Promega) according to the manufacturer’s instructions for plasmid expression or Lipofectamine 3000 (Invitrogen) for siRNA. siRNA sequences used were as follows: 5′-r(ACAAUCCUGAUCAGAAACC)d(TT)-3′ for nonspecific control siRNA, 5′-r(CUGGAAAGACUAGUGUGUU)d(TT)-3′ for human OPA1, 5′-r(GCAAUUAGCAGAGGCGUUA)d(TT)-3′ for human MIRO1, 5′-r(GCGUGGAGUGUUCGGCCAA)d(TT)-3′ for human MIRO2, 5′-r(GGGAAUUUGUGAGACAGUA)d(TT)-3’ for human G3BP1, and 5’-r(GGAACAAGAAGAAAGACAACC)d(TT)-3’ for human G3BP2.

### Live-cell and fixed cell confocal imaging

Live-cell imaging was performed using a Nikon Ti2-E–inverted fluorescence microscope equipped with a CSU-W1 spinning disk confocal scanner (Yokogawa) and an iXon Life 888 EMCCD camera (Andor) using a 100x/1.49 NA oil objective. Cells were maintained in phenol-red–free DMEM (Gibco) supplemented with 100 μM sodium pyruvate (Gibco) and 1% penicillin/streptomycin (Gibco) at 37°C in a humidified chamber with 5% CO_2_. Time-lapse images were acquired over a 10 min period using a cycle of 1-min imaging at 2-s interval and 1-min rest to minimize phototoxicity.

For fixed-cell imaging, overexpressing and immunostained samples were prepared. Cells were fixed with 4% paraformaldehyde (PFA; Biosesang) in phosphate-buffered saline (PBS; Enzynomics) for 10 min. Samples for immunostaining were further permeabilized with 0.5% Triton X-100 (Sigma-Aldrich) in PBS for 10 min and blocked with 1% bovine serum albumin (BSA; VWR) in PBS for 1 hour at room temperature. Next, samples were incubated with primary antibodies overnight at 4°C, followed by secondary antibody incubation for 1 hour at room temperature. All images were acquired using the same spinning disk confocal system. Z-stack images were collected for three-dimensional reconstruction.

### Expansion microscopy

Expansion microscopy was performed according to previously described ChromExM protocols (Pownall et al, 2023). HeLa cells were seeded on a 12-mm cover glass (Marienfeld Superior) in a 24-well plate (Corning) and subjected to oxidative stress for 20 min. Cells were fixed with 3% PFA and 0.1% glutaraldehyde (Sigma-Aldrich), followed by post-fixation in 0.7% PFA and 1% acrylamide (Sigma-Aldrich). Fixed samples were subsequently processed through iterative gelation and expansion steps as previously described (Pownall et al, 2023). After multiple gelation stages, immunostaining was performed with anti-G3BP (Abcam) and anti-TOM20 (Proteintech) as the primary antibodies and Alexa Fluor 488-conjugated anti-rabbit IgG (Abcam) and Alexa Fluor 647-conjugated anti-mouse IgG (Abcam) as the secondary antibodies. To amplify the fluorescence signals, we performed a second round of immunostaining using anti-Alexa Fluor 647 (Immunology Consultant Laboratory) and anti-Alexa Fluor 488 (Invitrogen) as the primary antibodies and Alexa Fluor 488-conjugated anti-rabbit IgG (Abcam) and Alexa Fluor 647-conjugated anti-mouse IgG (Abcam) as the secondary antibodies. Expanded gels were trimmed into thin layers, sealed with Picodent Twinsil (Picodent), and imaged using spinning disk confocal microscopy within 24 hours.

### Dual-color dSTORM super-resolution microscopy

To detect single-molecule localization of stress granule components, fixed HeLa cells were permeabilized and immunolabeled with primary antibodies against anti-G3BP1 (Invitrogen), anti-RACK1 (Santa Cruz), and anti-HA (Previously Covance), followed by secondary labeling with Alexa Fluor 488-conjugated anti-rabbit IgG (Abcam) and Alexa Fluor 647-conjugated anti-mouse IgG (Abcam).

For imaging, the cells were embedded in OxEA buffer [50 mM β-mercaptoethylamine hydrochloride, 3% OxyFlour™ (Oxyrase), and 20% sodium DL-lactate solution (Sigma-Aldrich) in PBS with pH adjusted to 8.0–8.5]. We acquired dSTORM data from a custom-built microscope equipped with a Nikon microscope body Ti2-E with a 100x/1.49 NA oil objective and HILO (highly inclined and laminated optical) illumination. Two channels were acquired sequentially with a 30-ms exposure, with at least 30,000 frames collected per channel using an Andor iXon 888 EMCCD camera.

Raw data were processed with Fiji plugin ThunderSTORM (Ovesný et al, 2014) to reconstruct super-resolution images and extract fluorescence signal localizations. To ensure spatial accuracy, post-processing was performed using the cross-correlation method for drift correction. Stress granule clusters were identified from G3BP1 localizations using a Python-based algorithm adapted from the quantitative super-resolution (qSR) analysis framework (Andrews et al, 2018). The area of each granule was taken as the area of the convex hull of its constituent localizations, which corresponds to the granule cross-section within the illuminated section. We quantified the localizations of RACK1 or HA-mCherry-Raptor using custom-built MATLAB code to calculate the number of single-molecule signals within each of the defined stress granules. Graphs were generated by Origin.

### Power-law fitting for dSTORM data

The relationship between the projected area of a stress granule (*x*) and the number of apoptotic factor localizations it contained (*y*) was described by a power law,

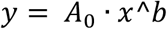

where *A*_0_ is the content predicted at *x* = 1μm^2^ and *b* is the scaling exponent. Residual dispersion increased with granule size on the linear scale, so both variables were log-transformed and the model was fitted by ordinary least squares as ln *y* = ln *A*_0_ + *b* ln *x*, implemented in Statsmodels (v0.14.4) in Python (v3.13.1). Granules with no detected localizations of the apoptotic factor were excluded from the log-scale fits (n = 56 of 2326 for RACK1; n = 3 of 1337 for Raptor). Repeating the analysis with all granules included using log(*N* + 1)gave essentially identical exponents, differing by at most 0.05 in any condition.

Localizations were fitted with a two-dimensional Gaussian PSF and no axial sectioning was applied at the analysis stage, so both the measured content and the measured area refer to the part of a granule falling within the HILO detection volume. Under constant internal concentration, the expected exponent follows from how a granule intersects that volume: a granule contained within it contributes its full content over its maximal cross-section, giving N ∝ A^1.5 for an approximately spherical granule, whereas a granule extending beyond it is sampled as a slab of approximately fixed thickness, giving N ∝ A. Exponents of 1.5 and 1 are therefore both reported as reference values, 1.5 being the more conservative, so that the comparison does not depend on the precise axial extent of detection volume. The fitted exponent was tested against each of these values with a two-sided *t* test on the regression coefficient. Exponents were compared between conditions by a two-sided z test on the difference of coefficients, with Holm correction across all pairwise comparisons.

Model comparison was performed on the log-transformed response, so that the power law, a linear model (y = a + bx) and an exponential model (y = a · exp(bx) + c) were all evaluated against the same quantity. For each model, predicted values μ(*x*) were computed and the residual sum of squares of ln *y* − ln μ(*x*) was minimized; the linear and exponential models were fitted on this scale by nonlinear least squares using the Levenberg-Marquardt algorithm in SciPy (v1.14.1). AICc was computed as 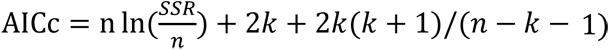, where SSR is the residual sum of squares, *n* the number of granules and *k* the number of fitted parameters, with *k* = 2 for the power-law and linear models and *k* = 3 for the exponential model.

### Proximity ligation assay (PLA)

PLA was performed using the Duolink In Situ Orange Starter Kit Mouse/Rabbit (Sigma-Aldrich) following the manufacturer’s protocol. HeLa cells were subjected to oxidative stress for 20, 40, or 60 min. Subsequently, cells were fixed, permeabilized, and incubated with anti-TOM20 (Proteintech) and anti-G3BP (Abcam). As a negative control, samples were incubated with anti-IgG antibody pairs (Sigma-Aldrich), followed by probe hybridization, ligation, and amplification reactions. PLA signals were acquired by spinning disk confocal microscopy, and z-stack images were collected for quantitative analysis.

### Monte Carlo simulation for stress granule–mitochondria proximity analysis

To assess the spatial relationship between stress granules and mitochondria, Monte Carlo simulations were performed. In each of 1,000 iterations per sample, stress granule positions were randomly repositioned within the cell mask, while the mitochondrial mask remained fixed. The mean distance between stress granule and mitochondria was calculated for each iteration, and the resulting simulated distribution was compared to the observed mean distance. Z-scores were calculated to quantify the deviation of the observed distance from the randomized mean. The simulation was implemented using Python (v3.13.1) with NumPy (v2.2.0) and SciPy (v1.14.1) libraries.

### Confocal image analysis

Fluorescence signals from PLA assays were analyzed using Imaris software. Z-stack images were reconstructed, and regions of interest corresponding to individual cells were traced according to DAPI nuclear staining. PLA signals within each cell were identified using the Spot plugin, and extracted metrics were exported for quantitative comparisons.

Stress granule number and size were quantified using Fiji (v1.52a, NIH). To improve signal clarity, at least 10 consecutive frames were averaged using Z-projection, followed by intensity thresholding and look-up table (LUT) adjustments to delineate stress granules from background. The “Analyze Particles” function was used to extract the total number and size of stress granules per cell.

Mitochondrial morphology and dynamics were analyzed using Fiji (v1.52a, NIH) and MitoMo2 (Zahedi et al, 2018). Mitochondrial structures were extracted using the “Skeletonize” plugin, and length metrics were computed using the “Analyze Skeleton” function. Time-lapse datasets were analyzed with MitoMo2 (Zahedi et al, 2018) to extract mitochondrial mobility indices and directional movement in each frame.

To evaluate spatial proximity between stress granules and mitochondria during stress responses, the shortest distance between the two organelles was calculated in cells fixed at 20, 40, and 60 min following arsenite treatment. After generating binary masks for stress granules and mitochondria in Fiji (v1.52a, NIH), the distances were quantified using a custom MATLAB script.

Cellular ATP levels were measured in HeLa cells expressing the genetically encoded ATP/ADP sensor PercevalHR. Cells were excited with 405 or 488 nm lasers, and emission was collected at 549 nm using spinning disk confocal microscopy. The PercevalHR fluorescence ratio (ATP excited at 488 nm / ADP excited at 405 nm) was quantified in Fiji (v1.52a, NIH) to evaluate cellular energy dynamics.

### Live-cell tracking analysis

Stress granule and mitochondrial dynamics were analyzed to assess their spatiotemporal associations. Fluorescence channels of time-lapse movies were separated using Fiji (v1.52a, NIH). Stress granules were segmented through Gaussian blur, LUT adjustments, and binary mask generation. Granule trajectories were extracted using the Fiji plugin TrackMate, which computed the x, y coordinates of individual granules across frames. Mitochondria were segmented by background subtraction and LUT adjustment, and their positional and dynamic features were quantified using Mitometer software (Lefebvre et al, 2021).

Tracking datasets were integrated using a custom MATLAB script to quantify three interaction metrics specifically for the pre-fusion segments of merging stress granule trajectories. The interorganelle distance was defined for each granule mask in each frame as the minimum Euclidean distance to nearest mitochondria mask. The metrics were: 1) stress granule fusion events, identified by merging of distinct trajectory segments in TrackMate; 2) proximity event rate, defined as the frequency of discrete encounter events, where each independent event is identified as an array of consecutive frames with an interorganelle distance of zero normalized to the total tracking duration of each granule; and 3) proximity occupancy, calculated as the total proportion of frames in which a stress granule was located at a distance of zero from mitochondria. A fusion event was classified as mitochondria-adjacent when the interorganelle distance of at least one of the two merging granules was zero in one or more frames of its pre-fusion segment; all other events were classified as non-mitochondria-adjacent.

Proximity occupancy sums all frames at zero interorganelle distance, whereas the event rate counts the discrete runs of such frames; the two metrics are therefore not independent and are reported as complementary descriptions of the same behavior.

### Cell recovery, kinetics, and viability assays

To assess cell recovery following stress, HeLa cells expressing mNeonGreen-G3BP1 and mCherry-TOMM20 were exposed to oxidative stress, and those with stress granules were randomly selected for tracking. After being transferred to the normal growth medium, the cells were tracked during 3-hour recovery period. At the end point, cells were classified as “recovered” if mNeonGreen-G3BP1 fluorescence became homogeneously cytoplasmic, indicating complete granule dissolution. Cells retaining visible granules or exhibiting apoptotic morphology (e.g., membrane blebbing) were categorized as “unrecovered.” For recovery kinetics analysis, cells were fixed at 0, 1, 2, and 3 hours after stress release. At each time point, randomly selected cells with stress granules were counted as unrecovered, whereas those without observable granules were counted as recovered. Percentage of recovered cells was calculated and compared across time points.

Cell viability was assessed by flow cytometry using the Annexin V-FITC Apoptosis Detection Kit (Abcam) according to the manufacturer’s instructions. Cells treated with DMSO or oligomycin A were subjected to oxidative stress, stained, and analyzed using S3e Cell Sorter (Bio-Rad) and FlowJo (v10.10.0).

### Mitochondrial membrane potential (MMP) measurement

To measure MMP, HeLa cells were treated with culture medium containing 100 nM TMRE (Abcam) for 15 min at 37°C with 5% CO_2_. After staining, the cells were washed with PBS to remove excess dye and minimize background fluorescence. TMRE fluorescence was detected using S3e Cell Sorter (Bio-Rad) and analyzed with FlowJo (v10.10.0).

### Immunoblotting

Cells were washed with ice-cold PBS, harvested, and lysed in 2× sample buffer (100 mM Tris-HCl adjusted to pH 6.8, 4% sodium dodecyl sulfate [SDS], 10% β-mercaptoethanol, 15% glycerol, and 0.008% bromophenol blue) at 95°C for 5 min. Lysates were separated by SDS–polyacrylamide gel electrophoresis and transferred to a nitrocellulose membrane (Amersham). The membranes were blocked for 1 hour at room temperature and then incubated with primary antibodies at 4°C overnight. After washing, HRP-conjugated secondary antibodies (Abcam) were applied at room temperature for 1 h. Protein bands were visualized using Amersham ImageQuant 800 (GE Healthcare) and quantified using Fiji (v1.52a, NIH).

### Statistical analysis

All data were obtained from at least three independently performed biological replicates unless stated otherwise. Graphs were generated using OriginPro 2019. Fluorescence images were analyzed using Fiji (v1.52a), Imaris, and custom MATLAB scripts. Live-cell tracking data were processed using the TrackMate plugin, Mitometer, and MitoMo2 software. Monte Carlo simulations were implemented using in-house Python code. Statistical analyses were performed using the two-tailed *t*-test for comparisons between two groups, one-way ANOVA with Tukey’s test for comparisons among multiple groups, Fisher’s exact test for comparisons of recovered versus unrecovered cell counts and the Mantel–Cox test for evaluating the cell recovery curves based on stress granule disassembly. Fitting and model comparison for the dSTORM scaling analysis are described separately above. Graphical data are presented as mean ± SEM unless stated otherwise. In the figures, significance follows: *P*-value ≥ 0.05 was considered as not significant (n.s.), \**P* < 0.05, \*\**P* < 0.01, \*\*\**P* < 0.001, and \*\*\*\**P* < 0.0001.

## Data availability

Computer code: Analysis code for live-cell tracking and statistical analyses. GitHub (https://github.com/wonkicholab/SG-Mito-Analysis). This study includes no data deposited in external repositories. All data are available in the main text or the supplementary materials. The expression plasmids reported in this manuscript are available upon request.

## Author contributions

**TLP:** Conceptualization; methodology; investigation; visualization; formal analysis; software; validation; writing – original draft; writing – review and editing. **GK:** Methodology; investigation; visualization; formal analysis; software; validation; writing – original draft; writing – review and editing. **HIK:** Methodology; investigation; visualization. **SD:** Investigation; visualization. **KR:** Software; validation. **YJL:** Validation. **CS:** Validation. **HS:** Methodology. **DKK:** Methodology. **YKK:** Methodology; writing – review and editing. **WKC:** Conceptualization; methodology; writing – review and editing; supervision; funding acquisition.

## Acknowledgements

We thank Dr. KJ Yoon for supporting materials. We thank Dr. ME Pownall for sharing detailed instructions of chromatin expansion microscopy and Dr. R Parker for helpful insights and discussion. We also thank the Stem Cell Center at Korea Advanced Institute of Science and Technology for assistance with flow cytometry and spinning disk confocal microscopy.

